# An Acyclic Phosphonate Prodrug of HPMPC is Effective Against VZV in Skin Organ Culture and Mice

**DOI:** 10.1101/2022.01.30.478368

**Authors:** M Lloyd, D Liu, J Lyu, J Fan, JM Overhulse, BA Kashemirov, MN Prichard, CE McKenna, JF Moffat

## Abstract

Varicella zoster virus (VZV) causes chicken pox and shingles and is prevalent worldwide. Acyclovir and penciclovir (and its prodrugs) are first-line treatments for VZV infections, but they are not highly potent against VZV and resistance may arise in immunocompromised people on long-term therapy. HPMPC (cidofovir) is active against VZV, but cidofovir is not approved for treating VZV diseases, is nephrotoxic, and is not orally bioavailable. Here, we present the synthesis and evaluation of USC-373, a phosphonate prodrug of HPMPC with activity against VZV and other DNA viruses. In cultured fibroblasts, it was potent against VZV Ellen laboratory strain and was not overtly toxic, with EC_50_ of 4 nM and CC_50_ of 0.20 μM, producing a selectivity index of 50. In ARPE-19 cells, USC-373 was effective against VZV-ORF57-Luc wild type strain and the acyclovir-resistant isogenic strain. In human skin organ culture, USC-373 formulated in cocoa butter and applied topically prevented VZV-ORF57-Luc spread without toxicity. In NuSkin mice with human skin xenografts, one daily dose of 3 mg/kg was effective by the subcutaneous route, and one daily dose of 10 mg/kg was effective by the oral route. Remarkably, a 10 mg/kg oral dose given every other day was also effective. USC-373 was well tolerated and mice did not lose weight or show signs of distress. The prodrug modifications of USC-373 increase the potency and oral bioavailability compared to its parent nucleoside analog, HPMPC.

## 1. Introduction

Varicella zoster virus (VZV) is a human-restricted alphaherpesvirus encoding 70+ genes required for replication, assembly, and virus spread (Cohen, 2010). VZV infections are ubiquitous worldwide, causing varicella (chicken pox) during primary infection and herpes zoster (shingles) during reactivation. The lifetime risk of zoster is approximately 30%, which corresponds to roughly 1 million cases in the U.S. per year (Yawn and Gilden, 2013). There are highly protective vaccines tailored to the populations most susceptible to varicella (children) and herpes zoster (adults over 50-60 years old) [reviewed in (Gershon and Gershon, 2013)]. Unfortunately, protection can wane and individuals, particularly immunocompromised patients that cannot receive the live-attenuated vaccine, may experience the severe, debilitating effects of a VZV infection. Antiviral drugs to treat VZV exist, however they must be taken within three days of onset and some have side effects (Sauerbrei, 2016). Resistance to these antivirals does arise, although it is most common in immunocompromised patients (Morfin et al., 1999; Sauerbrei, 2016; Strasfeld and Chou, 2010). Thus, there is a need for new, alternative antivirals that are more potent and better tolerated than the current drugs, and for drugs that are effective against resistant VZV strains.

The antiviral drugs for VZV prevent viral replication by targeting the DNA polymerase, including acyclovir (ACV), ganciclovir (GCV), cidofovir (CDV) and foscarnet (Strasfeld and Chou, 2010). Brivudine (BVDU), a drug approved in Europe, also blocks the VZV DNA polymerase (De Clercq and Li, 2016). VZV thymidine kinase (TK) is a viral enzyme required for the first phosphorylation step of many antivirals, including brivudine, acyclovir and ganciclovir [reviewed in (De Clercq and Li, 2016; Strasfeld and Chou, 2010)]. Unfortunately, the most common mechanism of VZV antiviral resistance is through TK mutations, which occur at various nucleotide positions in the TK gene (Morfin et al., 1999; Strasfeld and Chou, 2010). In the event of a TK mutation, these antivirals do not go through the first phosphorylation step. The unphosphorylated forms of these antivirals are rendered inactive and cannot interfere with viral replication. Therefore, alternative antivirals are necessary to combat such antiviral resistance.

Recently, there has been a push to develop antiviral compounds that bypass steps in the phosphorylation process needed for conversion to the active form. Foscarnet and cidofovir do not require the VZV TK and instead are entirely phosphorylated by cellular enzymes. However, they are not considered first-line antiviral drugs for VZV (Strasfeld and Chou, 2010), as they are both associated with major side effects, particularly renal toxicity [reviewed in (Upadhyayula and Michaels, 2013)]. This highlights the need for antivirals that do not require viral TK for phosphorylation to their active form and do not cause severe side effects. A further drawback of foscarnet and cidofovir is their low bioavailability, which precludes oral administration.

Phosphonate antiviral prodrugs bypass the initial phosphorylation step in infected cells and are quickly converted to the active triphosphate form by cellular kinases. Their mechanism of action is incorporation of the triphosphate nucleoside analog into nascent viral DNA, causing chain termination. An *N*-C16 tyrosinamide phosphonate prodrug of HPMPA, USC-087, was recently shown in an animal model to be orally effective against human adenovirus, a DNA virus (Toth et al., 2018). Here, we present a novel *N*-C18 *cis*-9-alkenyl tyrosinamide phosphonate prodrug of HPMPC (Cidofovir), USC-373 (Figure 1) and evaluate its efficacy versus varicella zoster virus (VZV) in cell culture, skin organ culture and in a humanized mouse model using subcutaneous and oral delivery. In all cases, USC-373 exhibited potent antiviral effects with little to no deleterious effect on the model system. A key observation from this study was that treatment is effective even when delayed to 3 days post-infection and VZV replication is established. This is clinically relevant, as most antiviral treatment starts after the onset of symptoms, usually several days post-infection. We found that the USC-373 prodrug is highly effective orally against VZV and is well-tolerated, demonstrating its potential as a novel antiviral.

**Figure 1.**
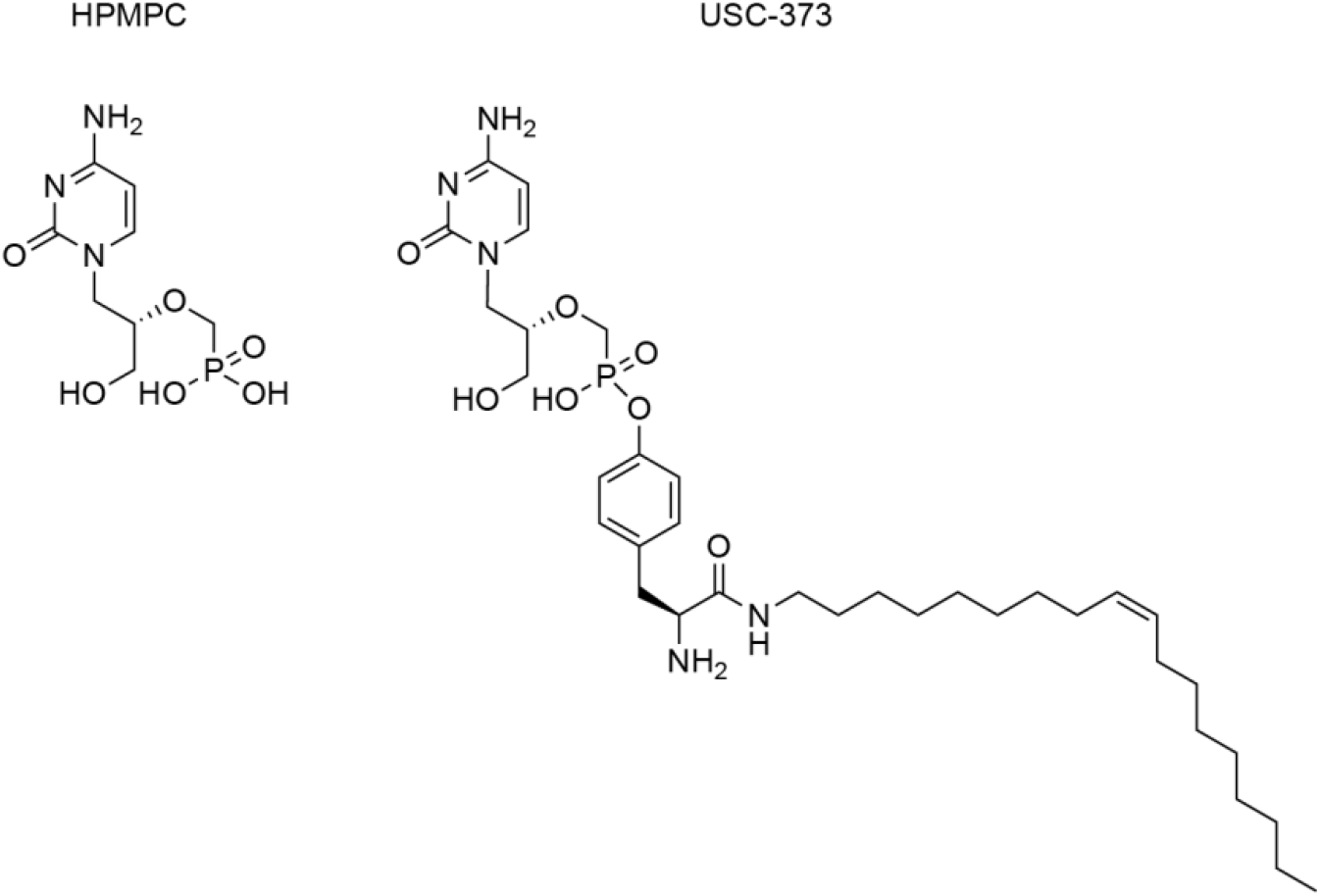
Chemical structures of USC-373 and HPMPC. USC-373 is an *N*-C_18_ *cis*-*9, 10* alkenyl tyrosinamide phosphonate ester prodrug of HPMPC.

## 2. Methods

### 2.1. Chemical synthesis of USC-373

USC-373 was prepared from HPMPC in three steps using the synthetic scheme presented below (Scheme 1). Intermediates and the product prodrug were characterized by LC/MS (LCQ Deca XP Max), ^1^H and ^31^P NMR (Varian VNMRS-500 NMR and Varian Mercury 400 2-channel NMR spectrometers). Final prodrug samples suitable for biological studies were purified by automated flash chromatography (ISCO Teledyne CombiFlash Rf+) on a silica gel column and further characterized by its UV/Vis spectrum (Beckman Coulter DU 800 spectrophotometer) and by combustion elemental analysis (Galbraith Laboratories).

#### (*2S*)-2-(*N*-(*t*-Butyloxycarbonyl)amino)-3-(4-hydroxyphenyl)-*N*’-(octadec-9-en-1-yl)propanamide, 1

The synthesis began with a reaction of commercially available *N*-Boc-L-tyrosine (2 mmol, Alfa Aesar, 98%) and oleylamine (2.4 mmol, Strem Chemicals, 95%) as a suspension in dichloromethane (DCM, 19 mL) with the addition of HOBt hydrate (2.6 mmol, Creosalus) and EDC-HCl (2.6 mmol, Chem-Impex International Inc). After stirring overnight at room temperature, the reaction mixture was checked for completion by TLC, diluted with 15 mL of DCM, and washed in series with 1.6 M citric acid (15 mL), saturated sodium bicarbonate (15 mL), and brine (15 mL, 3x). After evaporation, the crude product was purified by flash chromatography using a silica gel column eluted with ethyl acetate and hexanes to yield the tyrosine promoiety intermediate **1** as a white solid (87.9%, 0.933 g). ^1^H NMR (400 MHz, CD_3_OD): δ 7.08-7.06 (m, 2H, aromatic), δ 6.77-6.76 (m, 2H, aromatic), δ 5.39-5.35 (m, *J* = 14.0 Hz, 2H, C**H=**C**H**), δ 4.20-4.18 (m, 1H, C**H**NHBoc), δ 3.04-2.90 (m, 2H, NHC**H_2_**), δ 3.04-2.99 (dd, *J* = 6.0, 14.1 Hz, 1H, C**H_a_**H_b_(Tyr)), δ 2.95-2.90 (dd, *J* = 8.5, 14.0 Hz, 1H, CH_a_**H**_b_(Tyr)), δ 2.02-1.97 (m, 4H, C**H_2_**CH=CHC**H_2_**), 1.43 (s, 9H, NH**Boc**), δ 1.41-1.11 (m, 22H, 11CH_2_), δ 0.91-0.88 (m, 3H, **CH_3_**CH_2_).

#### Reaction of 1 with (*S*)-HPMPC to form protected prodrug derivative 2

To a suspension of (*S*)-HPMPC (1.07 mmol, APAC Pharmaceutical LLC, 99.90%) in dry DMF (13.3 mL), dry DIEA (20.4 mmol), tyrosine promoiety **1** (1.4 mmol), and (benzotriazol-1-yloxy)tripyrrolidino-phosphonium hexafluorophosphate (PyBOP) (3.33 mmol) were added. The reaction mixture was stirred under N_2_ at 40°C for 2 h and monitored by ^31^P NMR, with additional portions of PyBOP added as necessary to complete the reaction. DMF and DIEA were then removed under reduced pressure, and the residue was purified by flash chromatography to give cyclic prodrug intermediate **2** as a brown solid (38%, 0.314 g). ^31^P {^1^H} NMR (202 MHz, CD_3_OD) δ: 9.89 (s), δ 8.73 (s).

#### Deprotection of 2 to intermediate 3 and conversion to USC-373, 4

To a suspension of Boc-protected intermediate **2** (0.24 mmol) in DCM (5 mL) at 0°C, TFA (1.5 mL) was added dropwise. The reaction mixture was stirred at room temperature overnight. Volatiles were removed under reduced pressure, yielding the TFA salt of **3** as a brown oil. The crude product was used immediately for the next reaction. NH_4_OH (14.8 M, 18.2 mL) was added to 136.5 mL MeCN and the mixture was stirred for several minutes. Next, 80 mL of this NH_4_OH/MeCN solution was added to a round-bottom flask containing the cyclic (*S*)-HPMPC prodrug intermediate **3**. The mixture was shaken, swirled, and then sonicated until the prodrug dissolved, whereupon the remaining NH_4_OH/MeCN solution was added, and the reaction mixture was stirred at 45°C under N_2_ until all the starting material was consumed (^31^P, 10 h). Volatiles were then evaporated under reduced pressure and the residue was dissolved in MeOH. Cold DI water was added to the solution dropwise until a precipitate no longer formed. The solid was filtered and washed twice with cold DI water and dried. Purification by flash chromatography after dissolving in DCM gave the desired product **4** as a white solid (55%, 0.091 g). ^1^H NMR (400 MHz, CD_3_OD) δ: 7.64 (s, 1H), δ 7.62 (s, 1H) δ 7.18-7.12 (m, 2H, aromatic), δ 7.11-7.09 (m, 2H, aromatic), δ 5.81 (s, 1H, 5-H), δ 5.38-5.33 (m, *J* = 14.0 Hz, 2H, C**H=**C**H**), δ 4.06 (dd, *J* = 3.4, 13.9 Hz, 1H, C**H_a_**H_b_N), δ 3.98-3.94 (m, 1H, C**H**NH_2_), δ 3.83-3.77(m, 2H, CH_a_**H_b_**N, C**H_a_**H_b_P), δ 3.75-3.64 (m, 3H, C**H**OH, CH_a_**H_b_**P, C**H_a_**H_b_O), δ 3.52-3.48 (dd, *J* = 3.9, 12.1 Hz, 1H, CH_a_**H_b_**O), δ 3.23-3.19 (m, 2H, NHC**H_2_**), δ 3.15 (dd, *J* = 6.0, 14.1 Hz, 1H, C**H_a_**H_b_(Tyr)), δ 2.95-2.88 (dd, *J* = 8.5, 14.0 Hz, 1H, CH_a_**H_b_**(Tyr)), δ 2.07-1.94 (m, 4H, C**H_2_**CH=CHC**H_2_**), δ 1.51 (m, 2H, COCH_2_C**H_2_**), δ 1.41-1.11 (m, 22H, 11CH_2_), δ 0.91-0.88 (m, 3H, **CH_3_**CH_2_).; ^31^P {^1^H} NMR (202 MHz, CD_3_OD) δ: 13.49 (s).; MS m/z, calcd for C_35_H_58_N_5_O_7_P: 690.40. Found: 690.66. Combustion elemental analysis, % calcd for C_35_H_58_N_5_O_7_;P-2(H_2_O): C, 57.75; H, 8.59; N, 9.62. Found: C, 57.67; H, 8.22; N, 9.46.

Details about the purity analysis can be found in the supplemental file.

### 2.2. Propagation of cells and viruses

Human foreskin fibroblasts (HFFs) (CCD-1137Sk; American Type Culture Collection, Manassas, VA) were used prior to passage 20. Retinal pigment epithelial cells (ARPE-19; CRL-2302; ATCC) can be grown for 36+ passages. Cells were grown in Dulbecco’s Modified Eagle Medium (DMEM) with 4.5 g/L glucose, L-glutamine, and sodium pyruvate (DMEM 1X, Corning, Manassas, VA), and supplemented with 5-10% heat-inactivated fetal bovine serum (Benchmark FBS; Gemini Bio Products, West Sacramento, CA), penicillin–streptomycin (5000 IU/mL), and amphotericin B (250 μg/mL). The VZV strains included VZV-BAC-Luc (Zhang et al., 2007) and VZV-ORF57-Luc (Lloyd et al., 2020), both derived from the Parental Oka (POka, Accession number: AB097933) strain, VZV Ellen, a standard laboratory strain passaged more than 100 times since its isolation (Accession number: JQ972913.1), and acyclovir-resistant VZV (ACV^R^ VZV; (De et al., 2020)), derived from the VZV-BAC-Luc strain. VZV Ellen was propagated in HFFs, VZV-BAC-Luc and VZV-ORF57-Luc were propagated in both HFFs and ARPE-19 cells, and ACV^R^ VZV was propagated in ARPE-19 cells.

### 2.3. Preparation of human skin

Human fetal skin tissue (18-20 weeks gestational age, Advanced Bioscience Resources, Alameda, CA) was obtained adhering to all local, state, and federal guidelines. Adult human skin was obtained from reduction mammoplasty surgeries from healthy adults over 18 years old with informed consent and in accordance with approved Institutional Review Board protocols and procedures at SUNY Upstate Medical University in Syracuse, NY (SUNY Upstate, Institutional Review Board #1140572). Adult skin tissue was collected in saline within 2 h of surgery, and immediately processed for use as previously described (Haniffa et al., 2009; Lloyd et al., 2020). Briefly, whole thickness adult skin was cleaned, washed in sterile media, and thinned using a Weck knife and Goulian guard (0.028”) to approximately 700 microns. Fetal skin tissue was cleaned and washed in sterile media but did not require additionally thinning. Skin was cut into approximately 1-cm^2^ pieces and cultured on NetWells for skin organ culture (Corning, NY) or implanted into mice.

### 2.4. Compounds and formulations

All compounds were prepared from a dry powder at 10 mg/mL in dimethyl sulfoxide (DMSO) and stored at −20°C. Compounds used include USC-373, HPMPC, and acyclovir (ACV). The control compounds HPMPC (Cidofovir; BEI Resources, Manassas, VA) and ACV (Millipore Sigma, Burlington, MA) are commercially available. For cell-based assays, stock compounds were diluted in complete tissue culture medium. For all skin organ culture experiments, dry compounds were mixed with melted cocoa butter and vortexed to dissolve. If necessary, the mixture was sonicated briefly to break up large chunks of compound. The cocoa butter formulations were hardened on ice, then used at room temperature. For subcutaneous and oral delivery in mouse experiments, USC-373 was diluted in DMSO, then mixed 1:1 with Cremophor^®^ EL (Sigma C-5135; Millipore Sigma, Burlington, MA) and 8 parts normal saline (CDS solution) and stored at 4°C. For mouse experiments, HPMPC was prepared in water at 3 mg/mL and stored at 4°C.

### 2.5. Efficacy and cytotoxicity in cultured cells

Antiviral activity against VZV Ellen was evaluated in cytopathic effect (CPE) reduction assays by standard methods established previously (Prichard et al., 2006; Prichard et al., 2009). Briefly, monolayers of primary HFF cells were infected with virus at a multiplicity of infection of approximately 0.01 plaque forming units per cell, and CPE was evaluated 14 d post-infection. Concurrent cytotoxicity studies were performed using the same cells and compound exposure, and cell number was determined using CellTiter-Glo (Promega). Data obtained were used to calculate concentrations of compounds sufficient to inhibit viral replication by 50% (EC_50_) and cell number 50% (CC_50_). Multiple assays were performed for each compound to obtain statistical data.

### 2.6. Skin organ culture

Skin pieces were cultured on NetWells designed to hold tissue at the air-liquid interface and incubated at 35°C (adult) or 37°C (fetal) in a humidified CO_2_ incubator. VZV-BAC-Luc or VZV-ORF57-Luc were grown in HFFs and ARPE-19 cells to >75% CPE. Infected cells were harvested with trypsin/EDTA and sonicated to release virions and create a cell-free virus inoculum. Skin was inoculated with VZV (1×10^4^-10^5^ pfu/mL; 30 μL inoculum) by scarification with a 27-gauge needle (Lloyd et al., 2020; Rowe et al., 2010; Taylor and Moffat, 2005). Two hours after inoculation, skin was returned to NetWells and compounds or vehicle dissolved in cocoa butter were applied to the epidermis. For bioluminescence imaging, skin explants were submerged in D-luciferin (300 μg/mL in PBS) for 40 min before scanning in the IVIS™ 50. The medium (complete DMEM with 4-10% FBS) containing test compounds or vehicle was refreshed every other day after bioluminescence imaging. At termination of the experiment, skin was collected and fixed in 4% paraformaldehyde and stored at 4°C for histopathological analysis.

### 2.7. Histopathological Analysis

Skin pieces were fixed in paraformaldehyde and sent to HistoWiz Inc. (histowiz.com, New York, NY) for processing by a standard operating procedure and fully automated workflow. Samples were processed, embedded in paraffin, and sectioned at 5 μm. Hematoxylin and eosin (H&E) staining was performed per standard operating procedure. After staining, sections were dehydrated and film coverslipped using a TissueTek-Prisma and Coverslipper (Sakura). Whole slide scanning (40x) was performed on an Aperio AT2 (Leica Biosystems).

### 2.8. Animal procedures

Human adult skin xenografts were introduced subcutaneously above the left flank of 5– 6-week-old, male and female athymic nude mice [CR ATH Ho; Crl:NU(NCr)-*Foxn1^nu^*, Charles River, Wilmington, MA]. Single, full thickness xenografts were placed as previously described (Lloyd et al., 2020; Moffat et al., 1995). Three to four weeks after implantation, xenografts were inoculated with VZV-ORF57-Luc (cell-associated in HFFs, 1×10^6^ pfu/mL, 60 μL inoculum) by intradermal injection and scarification (Lloyd et al., 2020; Rowe et al., 2010). Virus growth was monitored daily using the IVIS™ 200 for bioluminescence imaging. Test compounds in CDS were administered by subcutaneous injection or oral gavage. Mice were weighed daily and observed for signs of distress. All animal procedures were performed with approved protocols under the guidance of the Committee for Humane Use of Animals at SUNY Upstate Medical University.

### 2.9 Bioluminescence imaging

Imaging was performed as described in (Rowe et al., 2010). Skin organ cultures were scanned with the IVIS™ 50 instrument and images were acquired for 30 s – 1 min, depending on pixel saturation (Caliper Life Sciences/Xenogen, Hopkinton, MA). The extent of VZV infection was measured as Total Flux (photons/sec/cm^2^/steradian) in a region-of-interest (ROI) encompassing the skin piece. Mice were scanned with the IVIS™ 200 instrument and images were acquired for an initial exposure time of 5 min; if pixels were saturated, additional images with shorter exposure times were acquired. An ROI was drawn over the skin xenograft and the Total Flux was recorded. The fold change was calculated as the daily Total Flux divided by the lowest Total Flux value, usually 2 or 3 dpi.

### 2.10. Statistical analysis

The 50% effective concentration (EC_50_) and 50% cytotoxic concentration (CC_50_) values were calculated using two model systems, Yield-Density and Sigmoidal Models, by XLfit 5.3 software (ID Business Solution, www.idbs.comi). Other calculations and graphs were made using GraphPad Prism (Graph-Pad Software, San Diego, CA, www.graphpad.com). Data from mouse experiments were analyzed using one-way ANOVA and Dunnett’s Multiple Comparison post hoc test. A *p* ≤ 0.05 was considered statistically significant.

## 3. Results

### 3.1. USC-373 was effective against VZV in cultured cells

HPMPC is the parent molecule of USC-373 (Fig. 1), and has broad spectrum activity against DNA viruses, including VZV (De Clercq, 2019; Magee and Evans, 2012). It was not known whether USC-373 would have similar activity against VZV. The antiviral activity and cytotoxic effects of USC-373 was compared to HPMPC in cell culture assays. Antiviral activity was measured by HFF cell lysis due to VZV spread and reported as the 50% effective concentration, EC_50_ (Table 1). USC-373 was highly effective against VZV, with an EC_50_ value of 0.004 μM for USC-373, which is at least 100-fold more potent than HPMPC. In parallel assays at higher concentrations, compounds were evaluated in HFF monolayers for their 50% cytotoxic concentration, CC_50_ (Table 1). The CC_50_ was 0.20 μM for USC-373. These results generated a selective index (SI) of 50 for USC-373. Generally, the higher the SI value, the safer the compound, and a value over 10 is considered a good target compound. This prompted further evaluations in ARPE-19 cells, in human skin and in mice.

**Table 1.** Efficacy and cytotoxicity of USC-373 against VZV Ellen in HFF cells.

|  | VZV-Ellen<br>HFFs |  |  | VZV-BAC-Luc<br>ARPE-19 | VZV-ACV-R<br>ARPE-19 |
| --- | --- | --- | --- | --- | --- |
|  | EC<sub>50</sub> (μM) | CC<sub>50</sub> (μM) | SI* | EC<sub>50</sub> (μM) | EC<sub>50</sub> (μM) |
| USC-373 | 0.004 | 0.20 | 50 | --- | --- |
| HPMPC | 0.52 | 60 | 115.4 | 1.67 | 1.05 |
| Acyclovir | 4.10 | >150 | >36.6 | 2.53 | n.d. |

\*SI (selectivity index) calculated as CC<sub>50</sub> / EC<sub>50</sub>

Acyclovir (ACV) is a drug of choice for treating VZV infections, however ACV resistance has been reported clinically and can be a serious problem in immunocompromised patients (Piret and Boivin, 2016; Sauerbrei et al., 2011). Thus, it is important to evaluate the effects of new antiviral compounds against acyclovir resistant VZV (ACV^R^ VZV). Furthermore, acyclovir resistance is often mediated by mutations in the TK gene (De Clercq and Li, 2016; Strasfeld and Chou, 2010), which is required to phosphorylate acyclovir to its active form. The O-linkage present on this prodrug obviates the need for phosphorylation by TK. Acyclovir, HPMPC (Cidofovir), and USC-373 were evaluated against VZV-ORF57-Luc wild type and an isogenic ACV^R^ VZV variant in ARPE-19 cell monolayers (Fig. 2). As expected, HPMPC was effective against WT and ACV^R^ viruses at 10 μM. WT VZV was sensitive to acyclovir, while the ACV^R^ VZV was resistant. Notably, USC-373 was effective against both wild type and ACV^R^ VZV.

**Figure 2.**
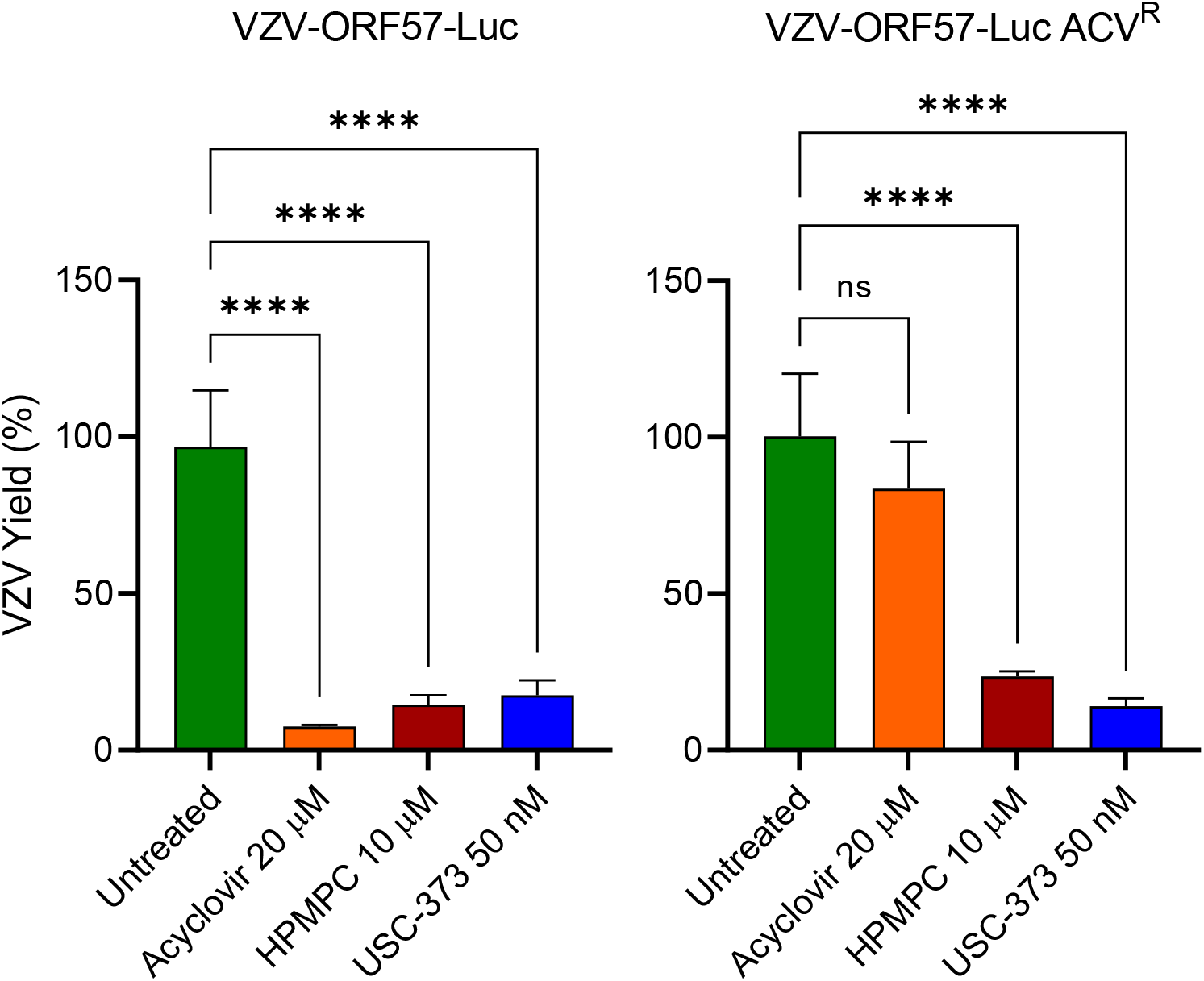
USC-373 is effective against ACV^R^ VZV in cultured cells. ARPE-19 cell monolayers were infected with VZV-ORF57-Luc wild type strain and the isogenic ACV^R^ variant (MOI = 0.01) for 2 h, then the virus was removed and replaced with tissue culture medium without antiviral compound (untreated), or with ACV 20 μM, HPMPC 10 μM, or USC-373 50 nM. VZV yield was measured by bioluminescence imaging after 2 days. Bars show the mean ± SD of 4 replicates. (****) signifies statistical significance between the treated and untreated groups (*p* < 0.0001, one-way ANOVA, Dunnett’s post hoc test), ns indicates “not significant”.

### 3.2. USC-373 prevents VZV spread in skin organ culture

The skin organ culture system is a secondary assay to evaluate antiviral compounds for efficacy against VZV. The full-thickness human skin explants provide a complex tissue microenvironment where antiviral compounds may have different activity than in cultured cells. It was not known whether the hydrophobic extensions on USC-373 would affect its activity in skin compared to HPMPC, which penetrates readily. Pieces of skin, approximately 1-cm^2^, were inoculated with VZV-ORF57-Luc by scarification and 2 h later were placed on NetWells above the medium at the air-liquid interface. Compounds dissolved in cocoa butter were added to the epidermal side. Groups included vehicle (cocoa butter alone), HPMPC 0.5%, and USC-373 (1.24%, equimolar to HPMPC). Treatments and tissue culture medium were refreshed every other day. After 5 days, the extent of VZV yield was measured by bioluminescence imaging and reported as Total Flux (proportional to pfu/mL and viral genome load (Zhang et al., 2007)). As expected, the cocoa butter vehicle slightly reduced the average Total Flux (Fig. 3). The process of adding and removing cocoa butter from the skin’s surface may dislodge VZV-infected cells, and so it has a consistent, minor effect on overall bioluminescence (data not shown). HPMPC and USC-373 reduced VZV yield in skin by approximately 1000-fold. HPMPC and USC-373 were also effective when added to the media (data not shown), however we were interested in topical formulations that could be applied directly to zoster lesions.

**Figure 3.**
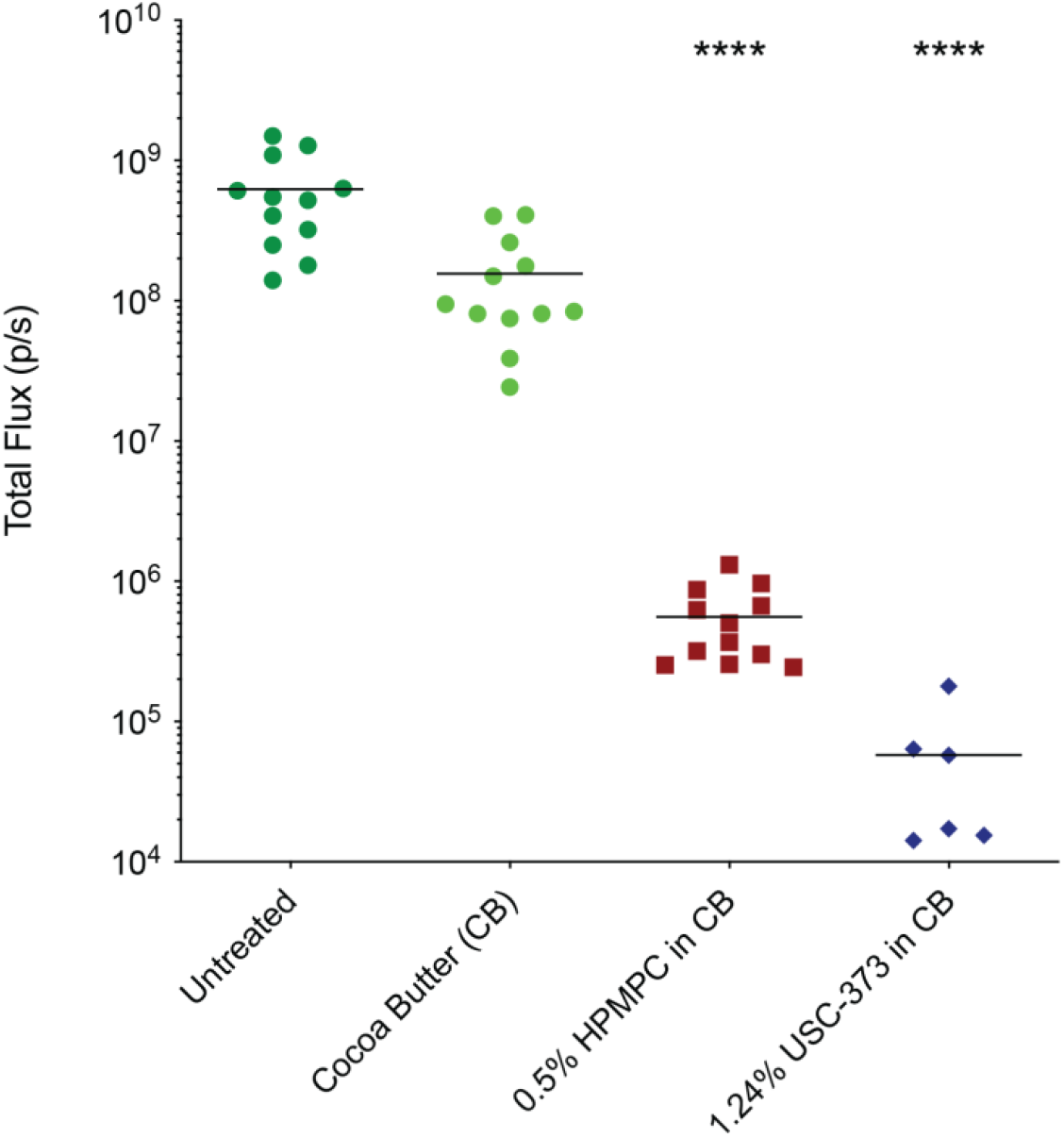
USC-373 prevents VZV spread in skin organ culture. VZV spread in skin organ culture was measured by bioluminescence imaging and measured as Total Flux (photons/sec/cm^2^/steradian) at 5 DPI. USC-373 was formulated in cocoa butter at equimolar weight-to-volume percentages compared to 0.5% HPMPC. The symbols represent individual pieces of VZV-infected skin (n=6-12); the black bar is the mean of the group. (****) signifies statistical significance between the treated groups and vehicle (cocoa butter) (*p* < 0.0001, one-way ANOVA, Dunnett’s post hoc test).

### 3.3. USC-373 was not toxic to skin when applied topically

While evaluating the efficacy of compounds in skin, it is also necessary to evaluate their toxicity. Fortunately, the bioluminescence signal at 5 dpi suggested that USC-373 and HPMPC were not overtly toxic to the skin, as luciferase activity is dependent on cell viability. Thus, we conducted histopathological analysis of skin after topical treatment. Similar to the skin organ culture experiments, 1-cm^2^ pieces of skin were inoculated with VZV by scarification, placed on NetWells, and treated with compounds in cocoa butter added to the epidermis. After 5 days, skin was fixed and processed for H&E staining (Fig. 4). In each panel, the skin was oriented with the epidermis at the top (stained purple) and the dermis at the bottom (stained pink). The untreated skin appeared normal after culture on NetWells, and it was used as baseline to evaluate the effects of each compound on the skin. The dermis was vascularized and highly cellular. The darker purple staining between the epidermis and dermis indicated the epidermal-dermal junction. Hair follicles were visible as invaginations of the epidermis into the dermis or large, purple, round to oval shapes in the dermis. No difference was seen between the untreated and vehicle only groups, with the epidermis present and epidermal-dermal junction fully intact. HPMPC-treated skin also appeared completely healthy. USC-373 was not toxic to the skin, as the epidermis was present and epidermal-dermal junction was intact. This supports the skin organ culture antiviral assay results.

**Figure 4.**
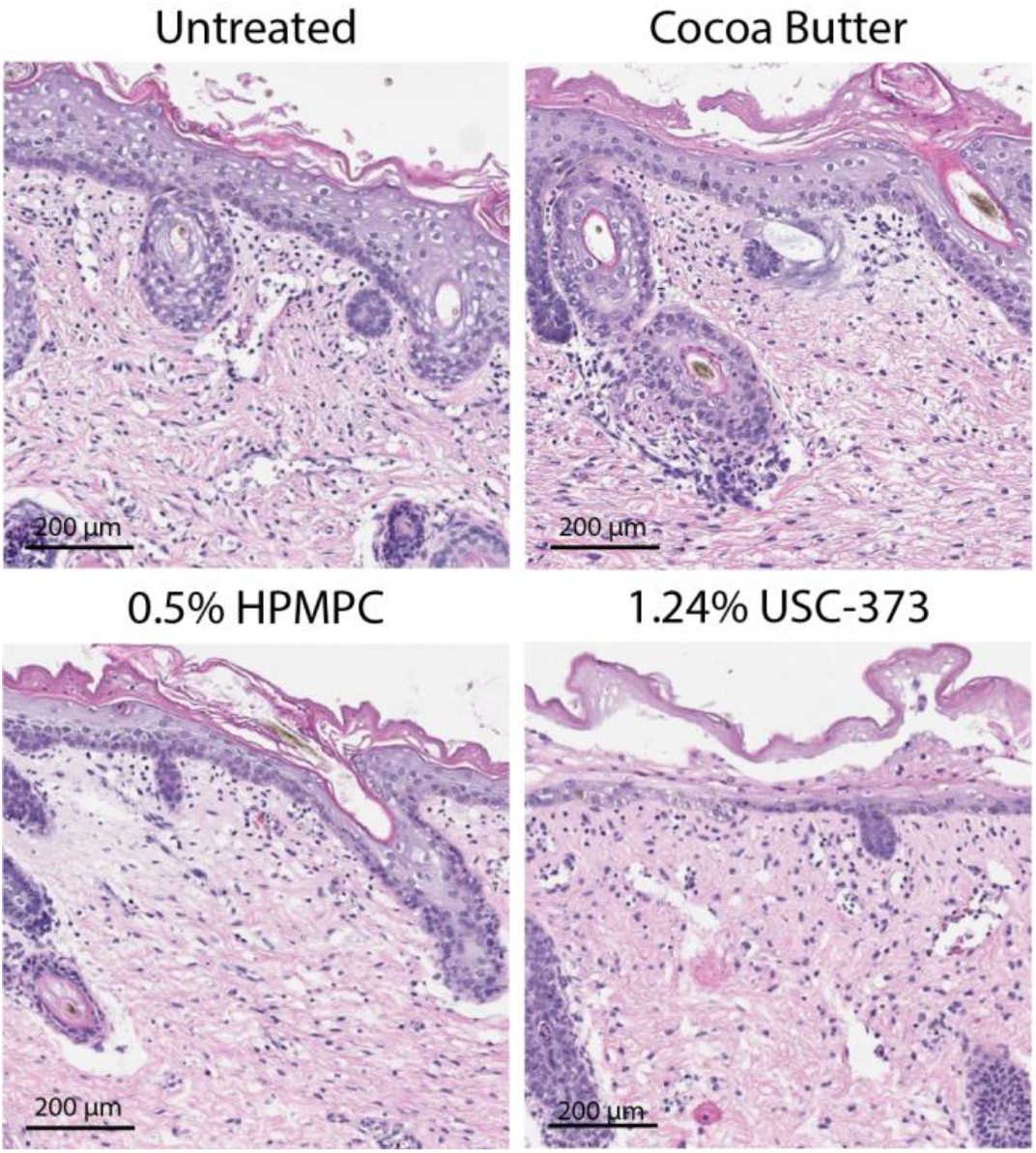
USC-373 is not toxic to skin when applied topically. Skin was infected with VZV and treated topically every other day with drugs formulated in cocoa butter at equimolar concentrations. Skin was collected 5 days post-inoculation (DPI) and fixed in 4% PFA for H&E staining. The skin is oriented with the epidermis at the top of the image (appears purple) and the dermis at the bottom (appears pink). Images are representative of tissues from multiple donors and experiments.

### 3.4. Efficacy of USC-373 in a humanized mouse model

It was not known whether USC-373 would be tolerated or if it would be bioavailable *in vivo* compared to HPMPC. NuSkin mice were inoculated with VZV by scarification and intraxenograft injection, and VZV spread was measured daily in the mice by bioluminescence imaging in the IVIS™ 200 instrument. The experimental groups included: vehicle, HPMPC, and USC-373. Different routes of administration (subcutaneous injection and oral gavage) were evaluated for the experimental compound; HPMPC must be administered by intraperitoneal (i.p.) injection. USC-373 was formulated in a Cremophor™-DMSO-saline solution (CDS) and administered according to the treatment schedules in Figs. 5A and 6A. VZV yield was calculated as the fold change in Total Flux after the last day of treatment. As expected, HPMPC was highly effective and significantly reduced VZV yield in all mouse studies (Fig. 5B and 6B). USC-373 significantly reduced VZV yield in a dose dependent manner. When given subcutaneously (Fig. 5), USC-373 prevented VZV spread at 10 mg/kg (p<0.001) and 3 mg/kg (p<0.01), but not at 1 mg/kg. When given orally (Fig. 6), USC-373 prevented VZV spread at 24.7 mg/kg (p<0.0001), which is equimolar to HPMPC 10 mg/kg, at 10 mg/kg (p<0.0001), and 10 mg/kg every other day (p<0.05). USC-373 was well-tolerated in the mice by both routes, and the mice did not lose weight or show signs of distress (Fig. 5C and 6C). Overall, USC-373 was 100x more potent than HPMPC and had the major advantage of oral bioavailability with little to no toxic effects *in vivo*.

**Figure 5.**
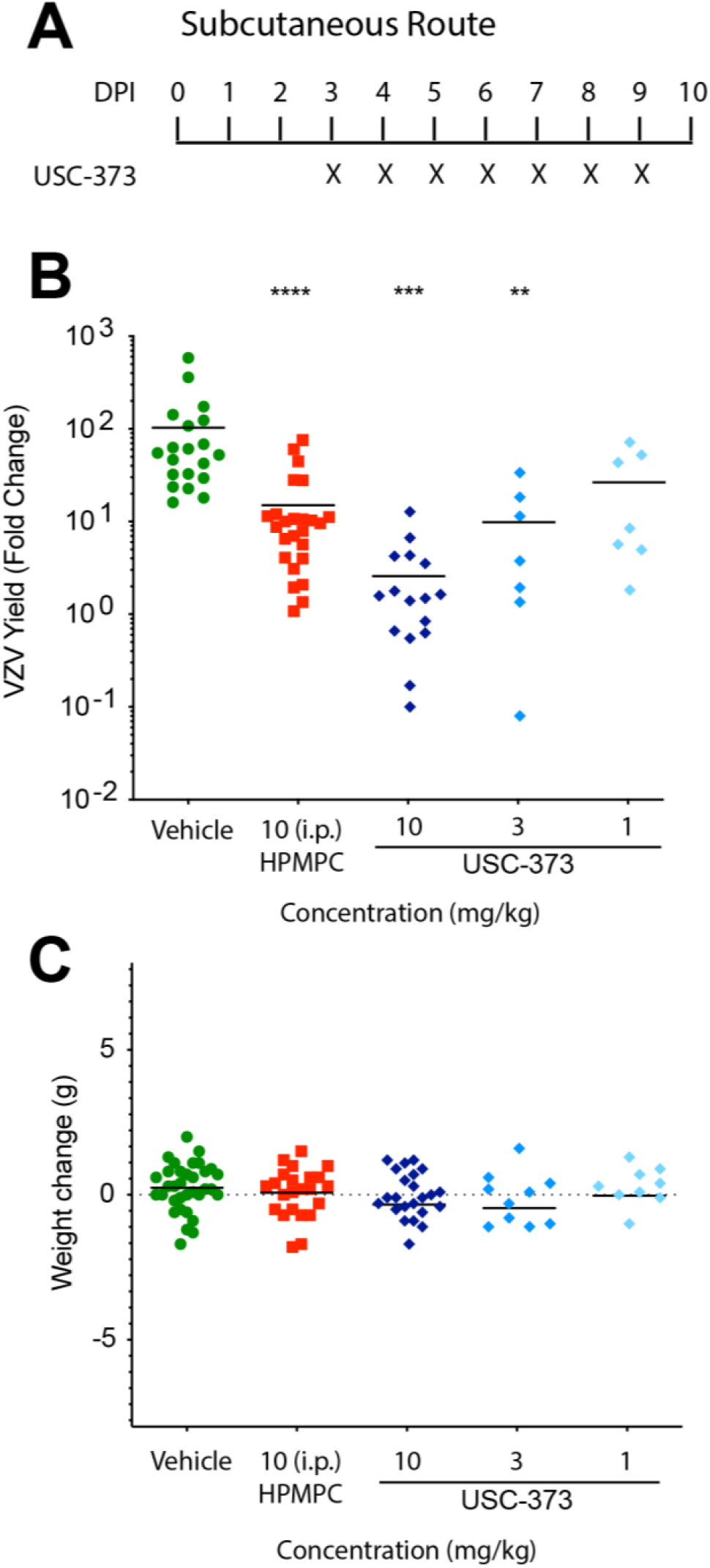
USC-373 prevents VZV spread and is well-tolerated *in vivo* when administered subcutaneously. Athymic nude mice with adult human skin xenografts were used to evaluate USC-373. The xenografts were inoculated with VZV by intraxenograft injection and scarification. (A) USC-373 was administered daily via the subcutaneous route, indicated by X. (B) Each symbol represents the fold change in VZV yield from a single mouse, calculated as the Total Flux on the day after the last treatment divided by the Total Flux on 2 or 3 DPI (the lowest value). (C) Each symbol represents the overall weight change (in grams) from the onset to conclusion of the study. The black bars represent the mean of each treatment group (n=9-20). These data are combined from multiple studies with multiple tissue donors. (*) signifies statistical significance between a treated group and vehicle (** < 0.01, *** < 0.001, one-way ANOVA, Dunnett’s post hoc test).

**Figure 6.**
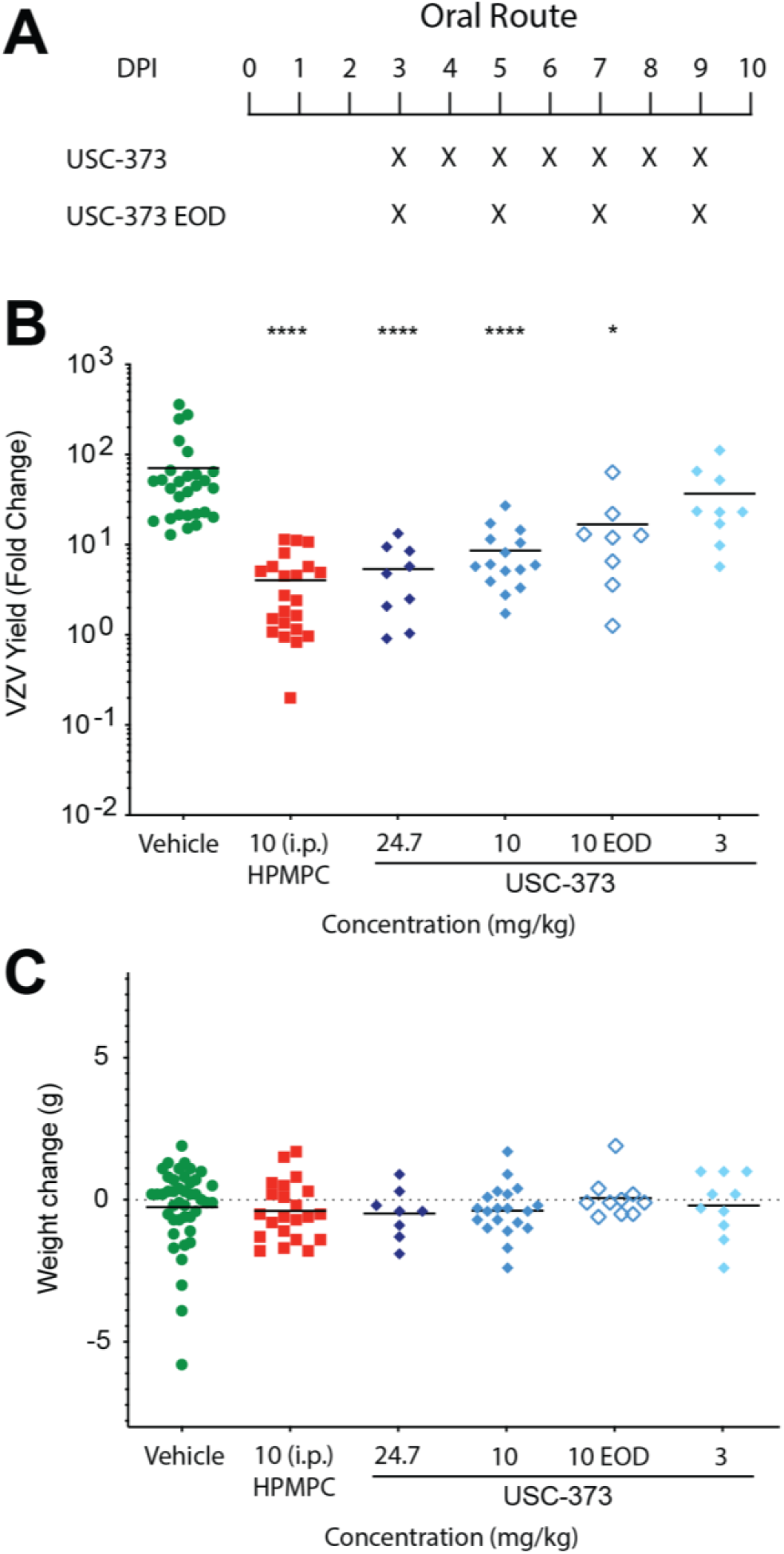
USC-373 prevents VZV spread and is well-tolerated *in vivo* when administered orally. Athymic nude mice with adult human skin xenografts were used to evaluate USC-373. The xenografts were inoculated with VZV by intraxenograft injection and scarification. (A) Compounds were administered daily via the oral route, indicated by X. (B) VZV yield, calculated as the Total Flux on the day after the last treatment divided by the Total Flux on 2 or 3 DPI (the lowest value). (C) Overall weight change, in grams, from the onset to conclusion of the study. Each symbol represents a single mouse. The black bar represents the mean of each treatment group (n=9-20). Data are combined from multiple studies and multiple tissue donors. (*) signifies statistical significance between a treated group and vehicle (* < 0.05, ** < 0.01, **** < 0.0001, one-way ANOVA, Dunnett’s post hoc test).

### 3.5. USC-373 prevents VZV rebound in vivo

Virus rebound after antiviral treatment is a serious issue and can lead to the development of resistance. The mice treated with USC-373, presented above, were evaluated for VZV rebound by bioluminescence imaging (Fig 7). As expected, VZV spread continued in the vehicle group up to DPI 14 in the first assay (Fig. 7A) and up to DPI 17 in the second assay (Fig. 7B). In the HPMPC group, VZV rebounded from DPI 10-14 or DPI 17-21, yet VZV yield remained significantly lower than the vehicle group (Fig. 7, *p* < 0.05, one-way ANOVA, Dunnett’s post hoc test). VZV rebound in the USC-373 groups was dose-dependent. The 3 mg/kg oral treatment did not suppress VZV spread after DPI 10. VZV rebounded somewhat from DPI 14-21 in the 10 mg/kg po EOD group. VZV was significantly suppressed in all other USC-373 groups (*p* < 0.05, one-way ANOVA, Dunnett’s post hoc test). The antiviral effects of USC-373, at a sufficient dose, prevented VZV spread *in vivo* during the treatment phase and suppressed rebound for up to 11 days.

**Figure 7.**
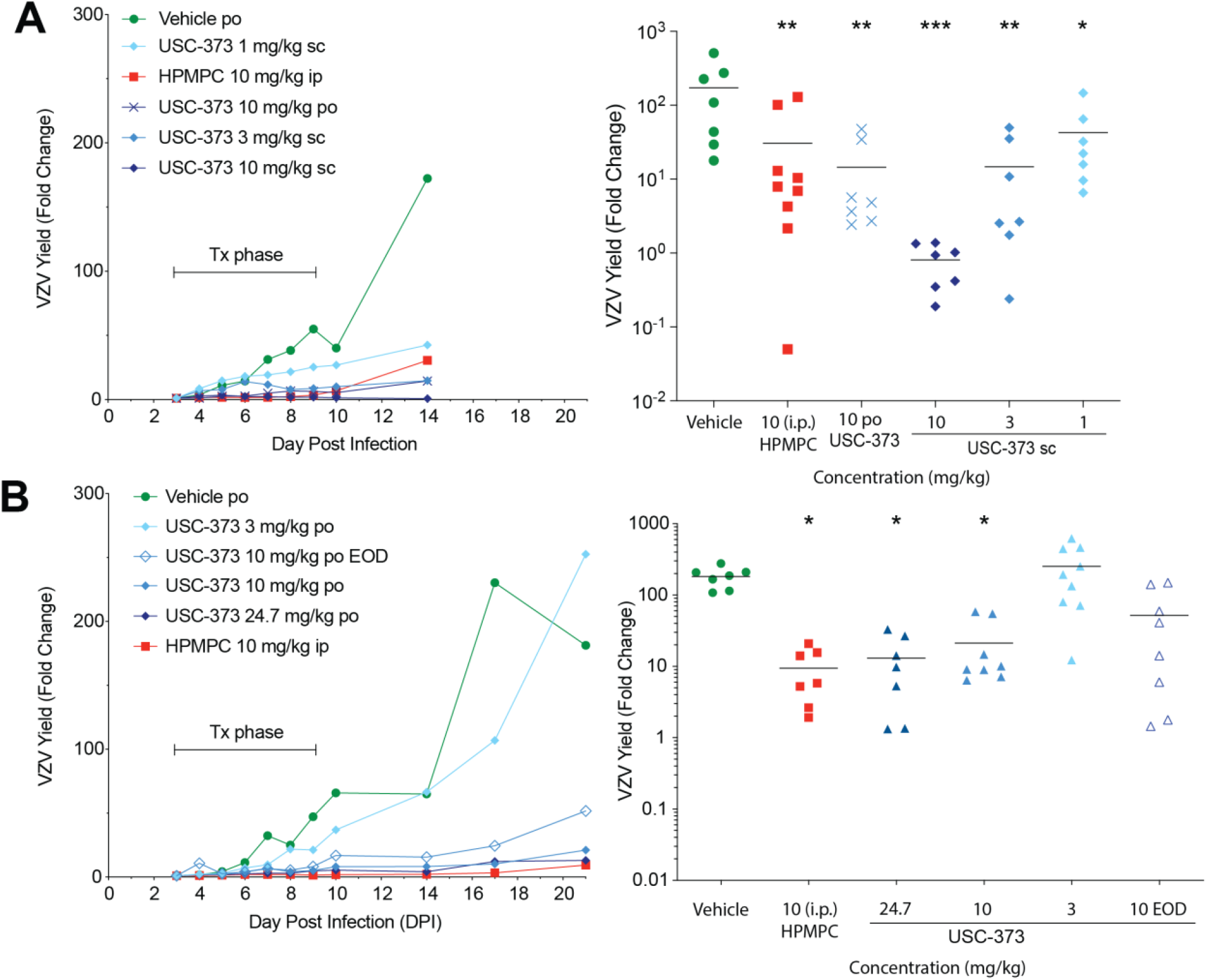
USC-373 prevents VZV rebound after treatment phase. Athymic nude mice with adult human skin xenografts were treated with USC-373 by subcutaneous or oral (po) routes, per the assays in Figs. 5 (shown in A) and 6 (shown in B). The rebound phase, following treatment from DPI 3-9, was measured by IVIS scans to DPI 14 (A) or DPI 21 (B). The right panels show the VZV yield at the end of the rebound phase compared to DPI 3. Each symbol represents one mouse with a line at the mean for the group, N=7-10. Each assay was conducted separately with different tissue donors. (*) signifies statistical significance between a treated group and vehicle (* *p* < 0.05, ** p < 0.01, *** p < 0.001, one-way ANOVA, Dunnett’s post hoc test).

## 4. Discussion

Here we present the synthesis and evaluation of a novel compound, USC-373, an *N*-C18 alkenyl tyrosinamide phosphonate ester prodrug of HPMPC, for activity against VZV. It was not known if USC-373 would be active against VZV. Indeed, we found that it prevented VZV spread in cell-based assays, skin organ culture, and humanized mice with skin xenografts.

USC-373 is an *N*-C18 cis-alkenyl tyrosinamide phosphonate ester prodrug of HPMPC that was designed to mimic the unsaturated fatty acid tail of the phospholipid bilayer. Lipid modifications are often utilized to enhance the PK properties of a drug to exhibit a longer circulation half-life by reabsorption across the tubular epithelium and increased serum protein binding. The chain length has a large impact on the various properties of the conjugate, with longer-chain fatty acids of >C14 generally showing increased lymphatic absorption and stability in the circulation when compared to shorter chain lengths. Oleic acid, a C18 alkenyl, is widely used as a liquid lipid in drug delivery due to its biodegradability and its biosafety (Sun et al., 2016). Studies have found that prodrug conjugates with unsaturated chains were more efficacious due to lower intracellular hydrolysis rates leading to longer intracellular retention (Zaro, 2015). Another study in 1987 by Jacob and coworkers (Jacob et al., 1987) also showed that with the introduction of unsaturation this resulted in an increase in bioactivity. We previously published the synthesis and antiviral evaluation of an *N*-alkylamido tyrosine ester of HPMPA, called USC-087, with a C16 alkyl chain (Toth et al., 2018). Through SAR studies, USC-373, containing this C18 alkenyl, was designed to mimic these lipid carriers that contain this oleic acid increasing its permeability and bioactivity.

The USC-373 compound studied here is a hundredfold more potent than the parent molecule, is active against a VZV strain that lacks thymidine kinase and has long-lasting effects *in vivo*. It is active in the low nanomolar range, and the selective index is acceptable. It is possible that this tool compound could be modified to reduce cytotoxicity. It will be important to address USC-373 effects on renal function because HPMPC (the drug cidofovir) is nephrotoxic (Izzedine et al., 2005). There is a need for additional antiviral therapies for herpes zoster that are more potent and can be given less often than the current drugs. The first-line antiviral drug is acyclovir and its derivative valaciclovir, the penciclovir derivative famciclovir, and brivudine in some countries (Andrei and Snoeck, 2021). In immunocompromised patients on long-term antiviral therapy, selection for virus mutants that are resistant to acyclovir typically occurs by mutation in ORF36, the TK gene. USC-373 does not require TK phosphorylation, due to its tyrosyl phosphonate linkage, and so it is effective against VZV strains that lack TK activity. It is likely that resistance would arise in ORF28, the DNA Pol catalytic subunit, but this has not yet been studied (Topalis et al., 2016). Resistance to antiviral drugs arises from a failure to suppress virus replication, so a major consideration should be developing new drugs with improved potency, selectivity, and long retention in the body (Griffiths, 2009).

USC-373 and HPMPC were effective topically, which opens avenues to develop potent antiviral treatments for herpes zoster. There are few options to treat herpes zoster lesions topically with direct acting antiviral drugs (Andrei and Snoeck, 2021). A 5% acyclovir cream is available for herpes labialis caused by HSV-1 and −2, but it is not effective for herpes zoster (De Clercq, 2004). Similarly, trifluridine and idoxuridine eye drops and creams are used to treat herpetic keratitis caused by HSV-1 and −2, but these have largely been replaced by oral antivirals for herpes zoster ophthalmicus (De Clercq, 2013; Johnson et al., 2015). Thus, there is an unmet need for direct acting antivirals that can be used in the eye and on the skin to treat herpes zoster. We found that USC-373 was well-tolerated on skin, and the hydrophobic chain may augment penetration into epithelial cells and keratinocytes. High potency and durability are also advantageous for developing a nonirritating and effective topical therapy for VZV infections. Future studies with skin explants and corneal epithelial cell cultures are warranted to address the optimal formulations, timing of application, and penetration of a compound such as USC-373 (Rönkkö et al., 2016).

The humanized mouse system is invaluable for studying VZV pathogenesis and evaluating antiviral compounds *in vivo* (Zerboni et al., 2014). VZV is a human restricted virus, and so humanized mouse models are required for such studies. The model we use has evolved from human fetal skin xenografts in SCID/*beige* mice (SCIDhu) (Rowe et al., 2010), to our current adult human skin xenografts in athymic nude mice (NuSkin) (Lloyd et al., 2020). The results presented in Figs. 5–7 were obtained using the NuSkin model, with skin tissue from multiple donors, which demonstrates the reproducibility of this system. We performed multiple in vivo assays and USC-373 was well tolerated in mice at all doses. Remarkably, USC-373 was effective orally at 10 mg/kg daily (eqimolar to 4 mg/kg HPMPC) as the usual positive control dose of 10 mg/kg HPMPC. Thus, USC-373 was superior to HPMPC in this study. A major advantage of USC-373 over HPMPC is the potent activity by the oral route. Another compound derived from HPMPC, brincidofovir (CMX001) is also orally bioavailable (Trost et al., 2015), suggesting that the alkyl moiety, or other similar molecular features, mediate absorption through the gastrointestinal tract. The USC-373 C16-alkyl side chain may also confer the durable suppression of VZV rebound that we observed for this compound (Fig. 7), which is a known effect of lipid modifications to prodrugs (Zaro, 2015).

The next steps for evaluating USC-373 are to isolate resistant VZV strains, to optimize the formulations for topical and oral dosing, and to address its pharmacology in the mouse and larger animals. These goals are beyond the scope of this initial report. A key question is whether USC-373 enters the brain, since VZV can cause meningoencephalitis, although rarely, and is linked to vasculitis and stroke (Nagel and Gilden, 2016). It is also important to address all aspects of dosing: formulations for optic, topical, and oral routes; the minimum effective dose; and dosing schedule. The formulation is a critical factor for hydrophobic compounds administered orally because the gastrointestinal tract has pH zones where absorption varies dramatically. This intriguing concept was described by Frenkel and colleagues who showed small aggregates, <100 nm, of non-nucleoside reverse transcriptase inhibitors (NNRTIs) were absorbed well in the GI tract (Frenkel et al., 2005). It is plausible that the ‘bent’ C18-alkenyl lipidomimetic modifier of USC-373 forms a lipid nanoparticle in aqueous solution, which may affect its absorption *in vivo*.

## Supplemental materials

Preparative chromatograms for **1**, **2**, and **4** with compound separation details; ^1^H, ^31^P and MS spectra.

## Acknowledgements

We dedicate this article to the memory of Mark Prichard, who was the first to demonstrate the high potency of USC-373.

CEM was supported by NIH grants R21 AI130927 and R01 AI135122 and JO was an NIH Chemical Biological Interface Trainee (T32 GM118289). JFM, MGL, and DL were supported in part by the contract HHSN272201700030I from the Division of Microbiology and Infectious Diseases, NIAID. We thank Inah Kang for assistance in preparing the manuscript.

## Scheme

**Scheme 1.**
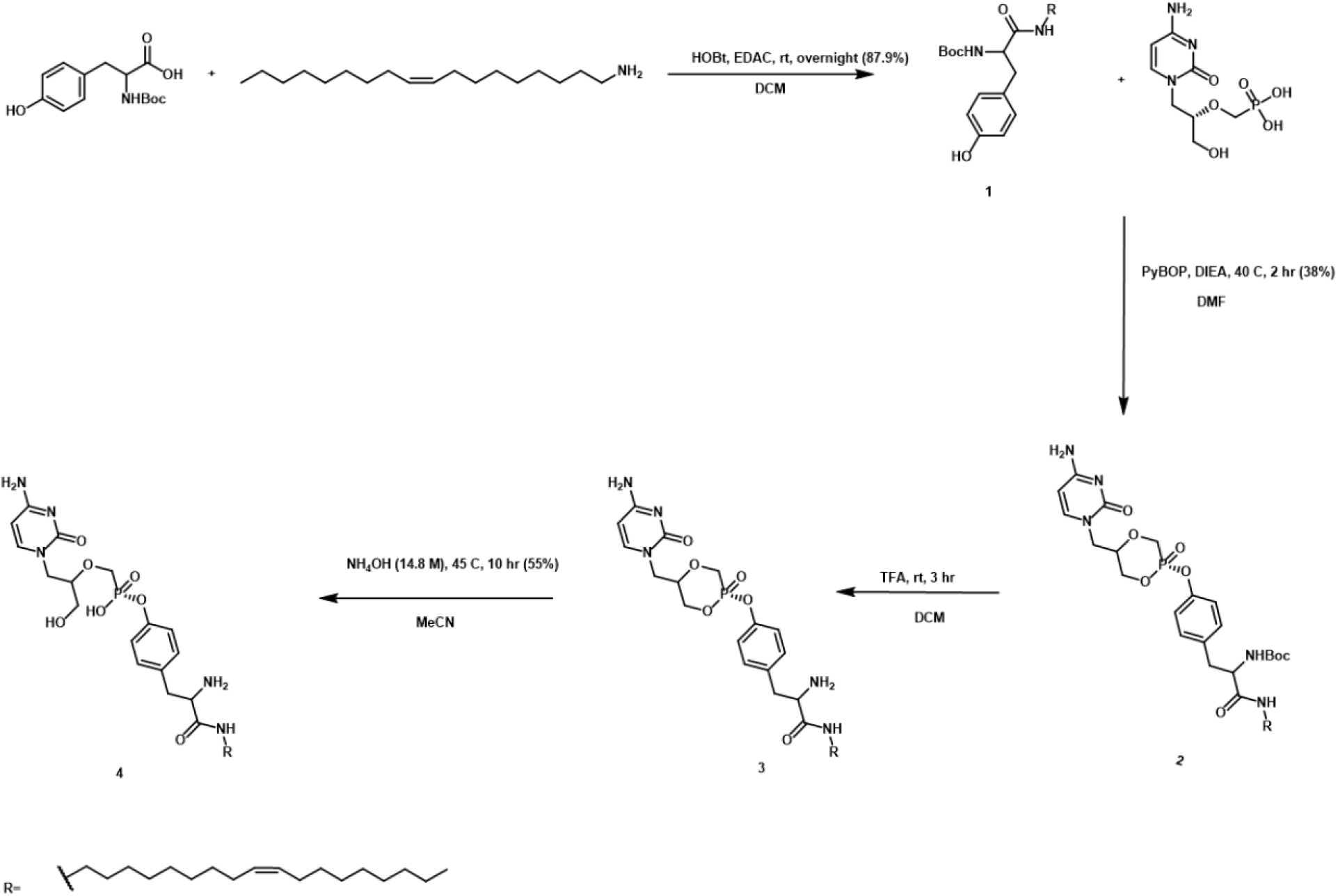
Synthesis of USC-373

## References

Andrei, G., Snoeck, R., 2021. Advances and Perspectives in the Management of Varicella-Zoster Virus Infections. Molecules 26.

Cohen, J.I., 2010. The varicella-zoster virus genome. Curr Top Microbiol Immunol 342, 1–14.

De, C., Liu, D., Depledge, D., Breuer, J., Singh, U.S., Hartline, C., Prichard, M.N., Chu, C.K., Moffat, J.F., 2020. β-L-1-[5-(E-2-Bromovinyl)-2-(hydroxymethyl)-1,3-dioxolan-4-yl)] uracil (L-BHDU) effectiveness against varicella-zoster virus and herpes simplex virus type 1 depends on thymidine kinase activity. bioRxiv, 2020.2002.2013.948190.

De Clercq, E., 2004. Antiviral drugs in current clinical use. Journal of clinical virology: the official publication of the Pan American Society for Clinical Virology 30, 115–133.

De Clercq, E., 2013. Selective anti-herpesvirus agents. Antiviral chemistry & chemotherapy 23, 93–101.

De Clercq, E., 2019. Fifty Years in Search of Selective Antiviral Drugs: Miniperspective. Journal of medicinal chemistry 62, 7322–7339.

De Clercq, E., Li, G., 2016. Approved Antiviral Drugs over the Past 50 Years. Clin Microbiol Rev 29, 695–747.

Frenkel, Y.V., Clark, A.D., Jr., Das, K., Wang, Y.H., Lewi, P.J., Janssen, P.A., Arnold, E., 2005. Concentration and pH dependent aggregation of hydrophobic drug molecules and relevance to oral bioavailability. Journal of medicinal chemistry 48, 1974–1983.

Gershon, A.A., Gershon, M.D., 2013. Pathogenesis and current approaches to control of varicella-zoster virus infections. Clin Microbiol Rev 26, 728–743.

Griffiths, P.D., 2009. A perspective on antiviral resistance. Journal of clinical virology: the official publication of the Pan American Society for Clinical Virology 46, 3–8.

Haniffa, M., Ginhoux, F., Wang, X.N., Bigley, V., Abel, M., Dimmick, I., Bullock, S., Grisotto, M., Booth, T., Taub, P., Hilkens, C., Merad, M., Collin, M., 2009. Differential rates of replacement of human dermal dendritic cells and macrophages during hematopoietic stem cell transplantation. J Exp Med 206, 371–385.

Izzedine, H., Launay-Vacher, V., Deray, G., 2005. Antiviral drug-induced nephrotoxicity. Am J Kidney Dis 45, 804–817.

Jacob, J.N., Hesse, G.W., Shashoua, V.E., 1987. .gamma.-Aminobutyric acid esters. 3. Synthesis, brain uptake, and pharmacological properties of C-18 glyceryl lipid esters of GABA with varying degree of unsaturation. Journal of medicinal chemistry 30, 1573–1576.

Johnson, J.L., Amzat, R., Martin, N., 2015. Herpes Zoster Ophthalmicus. Prim Care 42, 285–303.

Lloyd, M.G., Smith, N.A., Tighe, M., Travis, K.L., Liu, D., Upadhyaya, P.K., Kinchington, P.R., Chan, G.C., Moffat, J.F., 2020. A Novel Human Skin Tissue Model To Study Varicella-Zoster Virus and Human Cytomegalovirus. J Virol 94.

Magee, W.C., Evans, D.H., 2012. The antiviral activity and mechanism of action of (S)-[3-hydroxy-2-(phosphonomethoxy)propyl] (HPMP) nucleosides. Antiviral Res 96, 169–180.

Moffat, J.F., Stein, M.D., Kaneshima, H., Arvin, A.M., 1995. Tropism of varicella-zoster virus for human CD4+ and CD8+ T lymphocytes and epidermal cells in SCID-hu mice. J Virol 69, 5236–5242.

Morfin, F., Thouvenot, D., De Turenne-Tessier, M., Lina, B., Aymard, M., Ooka, T., 1999. Phenotypic and genetic characterization of thymidine kinase from clinical strains of varicella-zoster virus resistant to acyclovir. Antimicrob Agents Chemother 43, 2412–2416.

Nagel, M.A., Gilden, D., 2016. Developments in Varicella Zoster Virus Vasculopathy. Curr Neurol Neurosci Rep 16, 12.

Piret, J., Boivin, G., 2016. Antiviral resistance in herpes simplex virus and varicella-zoster virus infections: diagnosis and management. Curr Opin Infect Dis 29, 654–662.

Prichard, M.N., Keith, K.A., Quenelle, D.C., Kern, E.R., 2006. Activity and mechanism of action of N-methanocarbathymidine against herpesvirus and orthopoxvirus infections. Antimicrob Agents Chemother 50, 1336–1341.

Prichard, M.N., Quenelle, D.C., Hartline, C.B., Harden, E.A., Jefferson, G., Frederick, S.L., Daily, S.L., Whitley, R.J., Tiwari, K.N., Maddry, J.A., Secrist, J.A., 3rd, Kern, E.R., 2009. Inhibition of herpesvirus replication by 5-substituted 4’-thiopyrimidine nucleosides. Antimicrob Agents Chemother 53, 5251–5258.

Rönkkö, S., Vellonen, K.S., Järvinen, K., Toropainen, E., Urtti, A., 2016. Human corneal cell culture models for drug toxicity studies. Drug Deliv Transl Res 6, 660–675.

Rowe, J., Greenblatt, R.J., Liu, D., Moffat, J.F., 2010. Compounds that target host cell proteins prevent varicella-zoster virus replication in culture, ex vivo, and in SCID-Hu mice. Antiviral Res 86, 276–285.

Sauerbrei, A., 2016. Diagnosis, antiviral therapy, and prophylaxis of varicella-zoster virus infections. Eur J Clin Microbiol Infect Dis 35, 723–734.

Sauerbrei, A., Taut, J., Zell, R., Wutzler, P., 2011. Resistance testing of clinical varicella-zoster virus strains. Antiviral Res 90, 242–247.

Strasfeld, L., Chou, S., 2010. Antiviral drug resistance: mechanisms and clinical implications. Infect Dis Clin North Am 24, 809–833.

Sun, B., Luo, C., Li, L., Wang, M., Du, Y., Di, D., Zhang, D., Ren, G., Pan, X., Fu, Q., Sun, J., He, Z., 2016. Core-matched encapsulation of an oleate prodrug into nanostructured lipid carriers with high drug loading capability to facilitate the oral delivery of docetaxel. Colloids and Surfaces B: Biointerfaces 143, 47–55.

Taylor, S.L., Moffat, J.F., 2005. Replication of varicella-zoster virus in human skin organ culture. J Virol 79, 11501–11506.

Topalis, D., Gillemot, S., Snoeck, R., Andrei, G., 2016. Distribution and effects of amino acid changes in drug-resistant α and β herpesviruses DNA polymerase. Nucleic acids research 44, 9530–9554.

Toth, K., Spencer, J.F., Ying, B., Tollefson, A.E., Hartline, C.B., Richard, E.T., Fan, J., Lyu, J., Kashemirov, B.A., Harteg, C., Reyna, D., Lipka, E., Prichard, M.N., McKenna, C.E., Wold, W.S.M., 2018. USC-087 protects Syrian hamsters against lethal challenge with human species C adenoviruses. Antiviral Res 153, 1–9.

Trost, L.C., Rose, M.L., Khouri, J., Keilholz, L., Long, J., Godin, S.J., Foster, S.A., 2015. The efficacy and pharmacokinetics of brincidofovir for the treatment of lethal rabbitpox virus infection: a model of smallpox disease. Antiviral research 117, 115–121.

Upadhyayula, S., Michaels, M.G., 2013. Ganciclovir, Foscarnet, and Cidofovir: Antiviral Drugs Not Just for Cytomegalovirus. J Pediatric Infect Dis Soc 2, 286–290.

Yawn, B.P., Gilden, D., 2013. The global epidemiology of herpes zoster. Neurology 81, 928–930.

Zaro, J.L., 2015. Lipid-Based Drug Carriers for Prodrugs to Enhance Drug Delivery. The AAPS Journal 17, 83–92.

Zerboni, L., Sen, N., Oliver, S.L., Arvin, A.M., 2014. Molecular mechanisms of varicella zoster virus pathogenesis. Nat Rev Microbiol 12, 197–210.

Zhang, Z., Rowe, J., Wang, W., Sommer, M., Arvin, A., Moffat, J., Zhu, H., 2007. Genetic analysis of varicella-zoster virus ORF0 to ORF4 by use of a novel luciferase bacterial artificial chromosome system. J Virol 81, 9024–9033.

