## Supplemental Files for "An Acyclic Phosphonate Prodrug of HPMPC is Effective Against VZV in Skin Organ Culture and Mice"

Supplemental File:

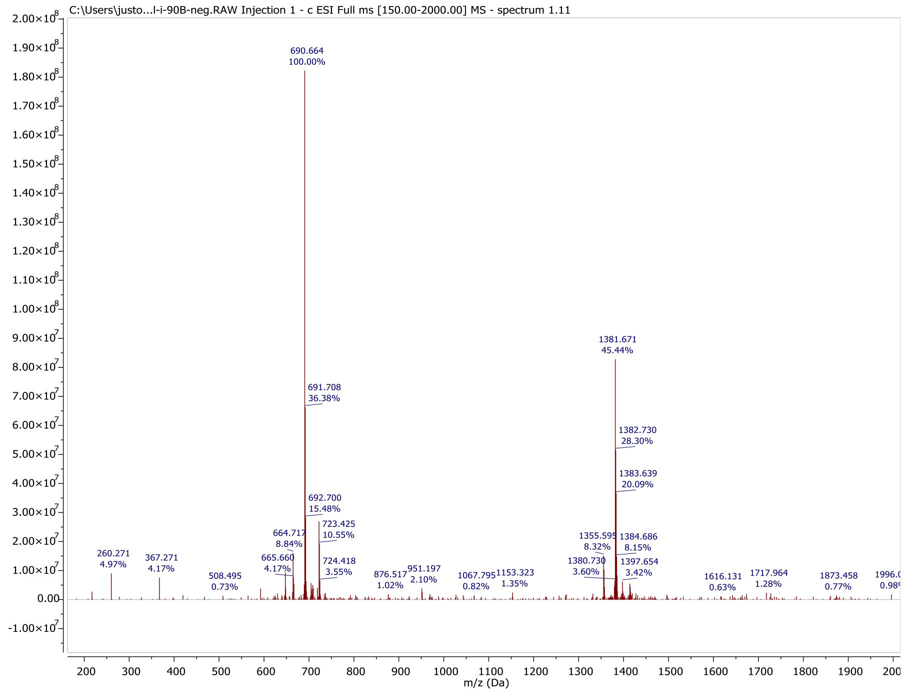


**Figure S1.** MS of USC-373 in negative mode. MS: *m/z* calcd ($C_{35}H_{57}N_5O_7P$) 690.40 (M-H)$^-$, found 690.664 (M-H)$^-$.

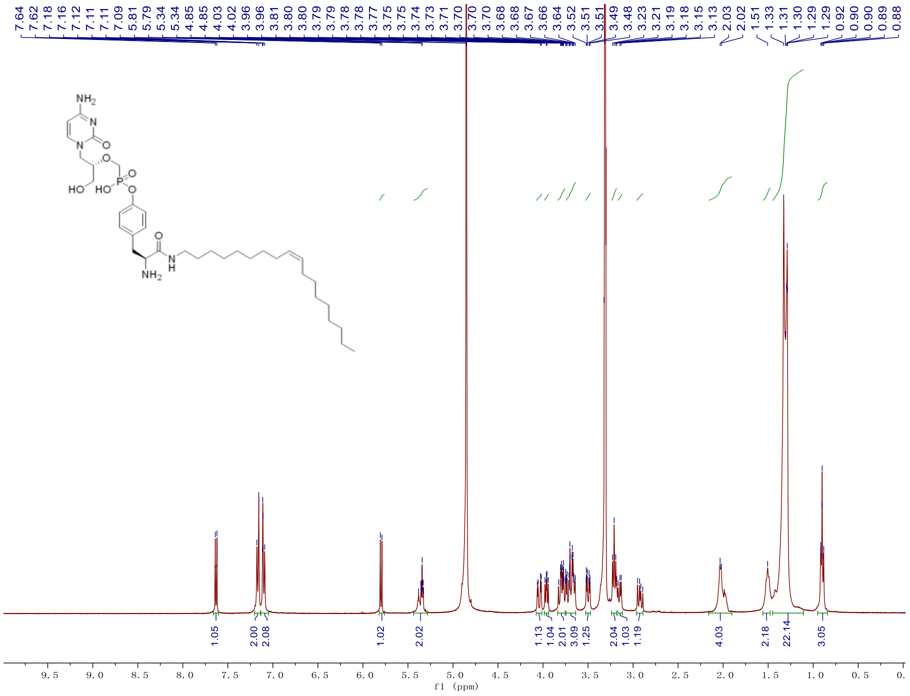

**Figure S2.** $^1$H NMR spectrum (400 MHz, $CD_3OD$) of USC-373: δ 7.64 (s, 1H), δ 7.62 (s, 1H) δ 7.18-7.12 (m, 2H, aromatic), δ 7.11-7.09 (m, 2H, aromatic), δ 5.81 (s, 1H, 5-H), δ 5.38-5.33 (m, *J* = 14.0 Hz, 2H, C**H=**C**H**), δ 4.06 (dd, *J* = 3.4, 13.9 Hz, 1H, C$\mathbf{H_a}H_b$N), δ 3.98-3.94 (m, 1H, C**H**$NH_2$), δ 3.83-3.77(m, 2H, $CH_a\mathbf{H_b}$N, C$\mathbf{H_a}H_b$P), δ 3.75-3.64 (m, 3H, C**H**OH, $CH_a\mathbf{H_b}$P, C$\mathbf{H_a}H_b$O), δ 3.52-3.48 (dd, *J* = 3.9, 12.1 Hz, 1H, $CH_a\mathbf{H_b}$O), δ 3.23-3.19 (m, 2H, NHC$\mathbf{H_2}$), δ 3.15 (dd, *J* = 6.0, 14.1 Hz, 1H, C$\mathbf{H_a}H_b$(Tyr)), δ 2.95-2.88 (dd, *J* = 8.5, 14.0 Hz, 1H, $CH_a\mathbf{H_b}$(Tyr)), δ 2.07-1.94 (m, 4H, C$\mathbf{H_2}$CH=CHC$\mathbf{H_2}$), δ 1.51 (m, 2H, $COCH_2$C$\mathbf{H_2}$), δ 1.41-1.11 (m, 22H, 11$CH_2$), δ 0.91-0.88 (m, 3H, $\mathbf{CH_3}CH_2$).

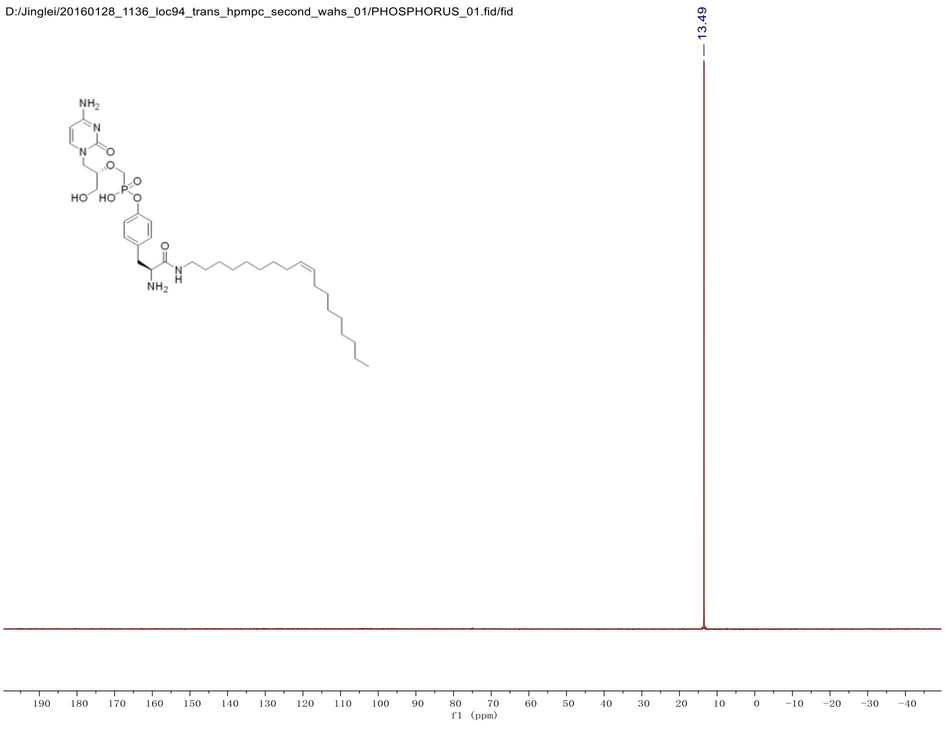


**Figure S3.** $^{31}$P {$^1$H} NMR spectrum (202 MHz, $CD_3OD$) of USC-373: δ 13.49 (s).

$$Dilution\ Factor * Abs = \varepsilon * \left(\frac{\frac{Mass\ of\ sample}{Molecular\ Weight}}{Stock\ Volume}\right)$$ Supplemental equation 1.

| **Stock Solution** | **3.19 mg USC-373 sample**<br>**Dissolved 1.020 mL EtOH** | | | |
|---|---|---|---|---|
| **Sample Preparation** | **49.8 µL stock solution + 2.040 mL EtOH** | | | |
| **Abs at 273 nm** | **0.8878** | **0.8569** | **0.9124** | **0.9019** |
| **Percent Content** | **93.7%** | **90.3%** | **96.3%** | **95.3%** |

**Table S1.** The average percent content of active compound USC-373 in sample by UV/Vis is 93.50% using supplemental equation 1. Concentration factor of this sample was 42. Abs at 273 nm was measured (Beckman Coulter DU 800 Spectrophotometer). Extinction coefficient for HPMPC, $\varepsilon$, used was 8362 at 274 nm in EtOH. Percent content was calculated subtracting 5% for the absorbance of tyrosine contribution in the sample.

| **Elemental Analysis** | **Nitrogen %** | **Carbon %** | **Hydrogen %** |
|---|---|---|---|
| **Galbraith Data** | 9.46 | 57.67 | 8.22 |
| **Calculated CHN of USC-373** | 10.12 | 60.76 | 8.45 |
| **Calculated CHN of USC-373 with 2 $H_2O$** | 9.62 | 57.75 | 8.59 |
| **Percent difference from original** | 0.16 | 0.08 | 0.37 |
| | **Molar Mass** | **Molar Mass Corrected** | **wt%** |
| | 691.85 | 727.88 | 95.1% |

**Table S2.** The percent content of active compound USC-373 in sample by elemental analysis (Galbraith Laboratories) is 95.1%. Molecular formula of USC-373: $C_{35}H_{58}N_5O_7P$. Molecular formula of USC-373 with 2 $H_2O$: $C_{35}H_{62}N_5O_9P$.

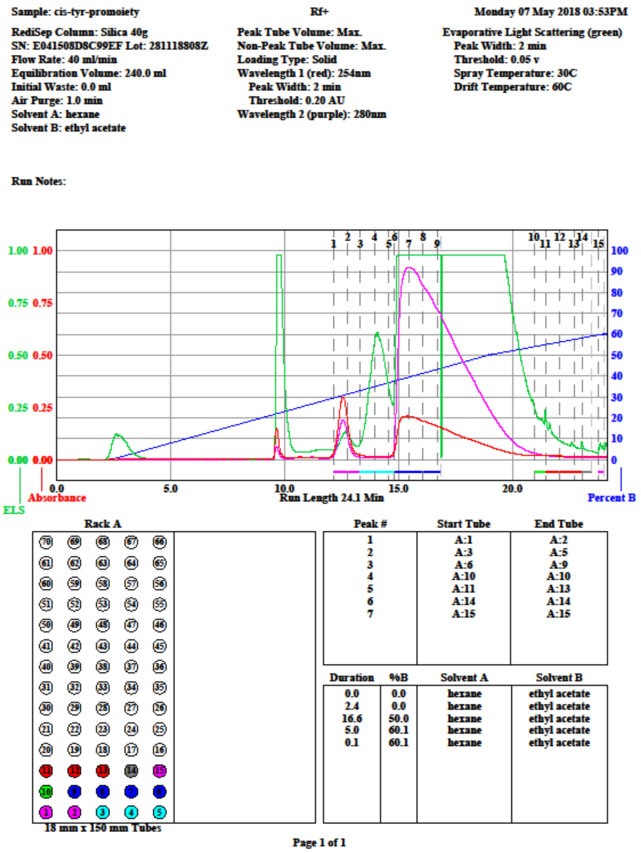


**Figure S4**. ISCO Teledyne CombiFlash Rf+ flash chromatography purification of tyrosine promoiety **1** with UV (254 nm:red trace; 280nm:purple trace) and evaporative light scattering detection (green trace). Promoiety eluted at 15 minutes.

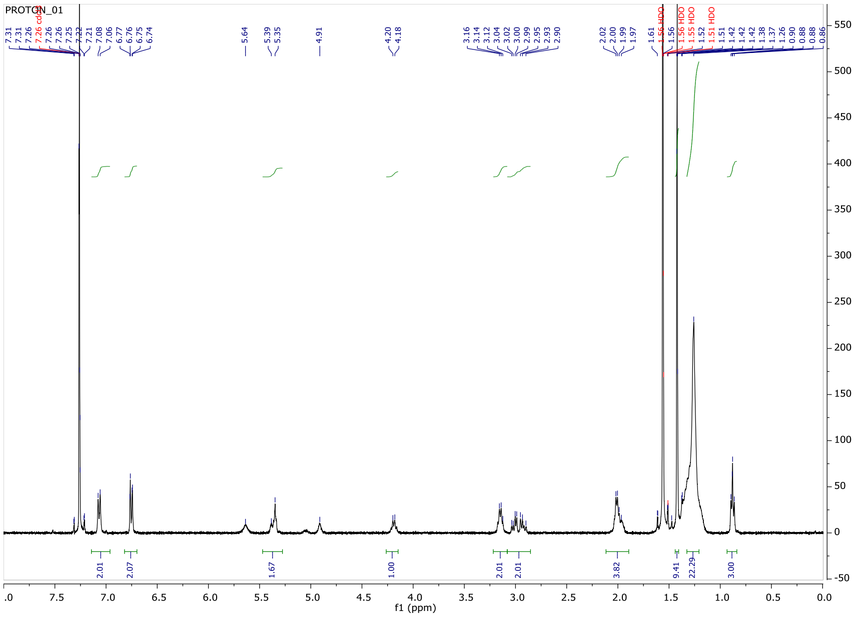


**Figure S5**. $^1$H NMR spectrum (400 MHz, $CD_3OD$) of purified tyrosine promoiety **1**: δ 7.08-7.06 (m, 2H, aromatic), δ 6.77-6.76 (m, 2H, aromatic), δ 5.39-5.35 (m, *J* = 14.0 Hz, 2H, C**H=**C**H**), δ 4.20-4.18 (m, 1H, C**H**NHBoc), δ 3.04-2.90 (m, 2H, NHC**H$_2$**), δ 3.04-2.99 (dd, *J* = 6.0, 14.1 Hz, 1H, C**H$_a$**H$_b$(Tyr)), δ 2.95-2.90 (dd, *J* = 8.5, 14.0 Hz, 1H, CH$_a$**H$_b$**(Tyr)), δ 2.02-1.97 (m, 4H, C**H$_2$**CH=CHC**H$_2$**), 1.43 (s, 9H, NH**Boc**), δ 1.41-1.11 (m, 22H, 11$CH_2$), δ 0.91-0.88 (m, 3H, **CH$_3$**$CH_2$).

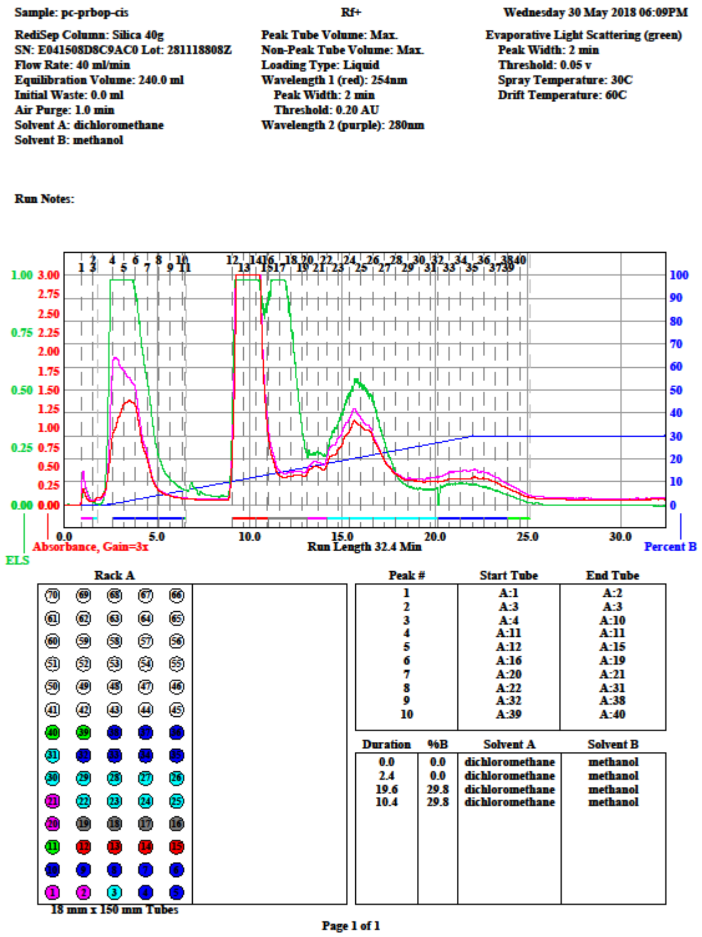


**Figure S6**. ISCO Teledyne CombiFlash Rf+ flash chromatography purification of intermediate compound **2** with UV (254 nm:red trace; 280nm:purple trace) and evaporative light scattering detection (green trace). Intermediate eluted at 15 minutes.

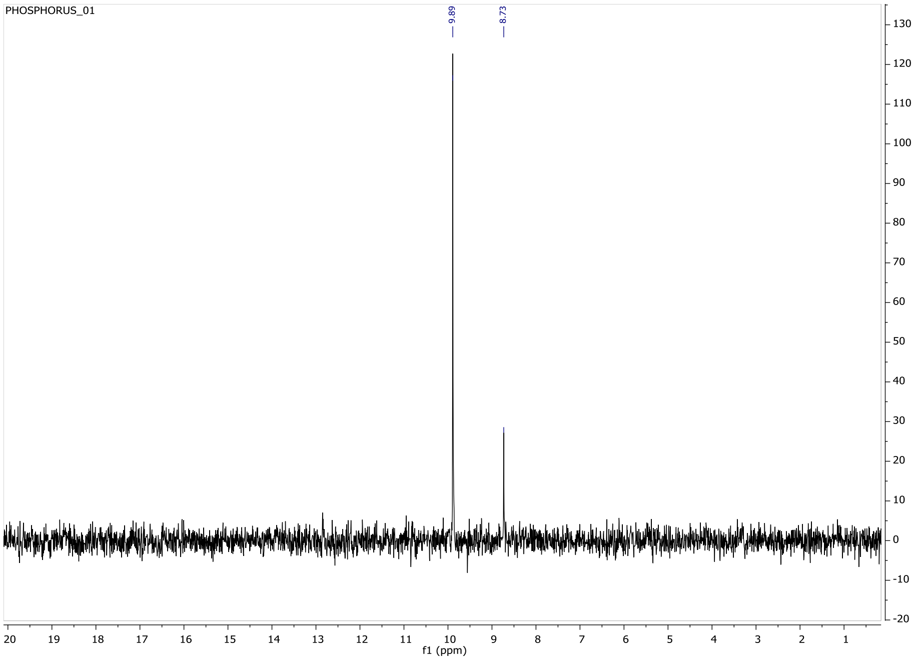


**Figure S7.** $^{31}P$ {$^{1}H$} NMR spectrum (202 MHz, $CD_3OD$) of purified **2**: δ 9.89 (s), δ 8.73 (s).

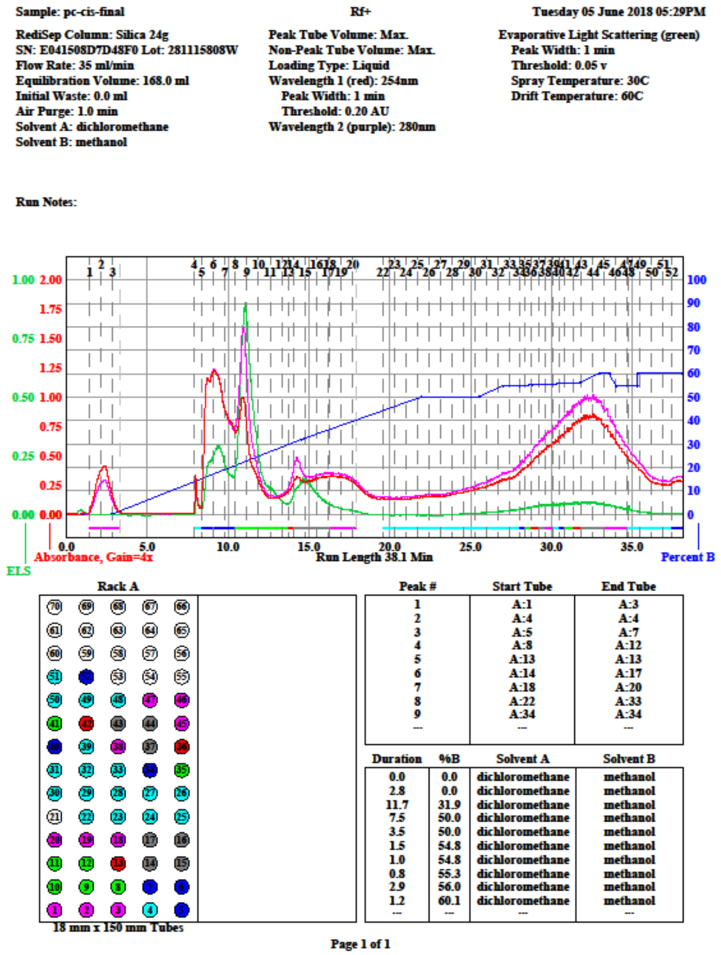

Sample: pc-cis-final

Rf+

Tuesday 05 June 2018 05:29PM

RediSep Column: Silica 24g
SN: E041508D7D48F0 Lot: 281115808W
Flow Rate: 35 ml/min
Equilibration Volume: 168.0 ml
Initial Waste: 0.0 ml
Air Purge: 1.0 min
Solvent A: dichloromethane
Solvent B: methanol

Peak Tube Volume: Max.
Non-Peak Tube Volume: Max.
Loading Type: Liquid
Wavelength 1 (red): 254nm
Peak Width: 1 min
Threshold: 0.20 AU
Wavelength 2 (purple): 280nm

Evaporative Light Scattering (green)
Peak Width: 1 min
Threshold: 0.05 v
Spray Temperature: 30C
Drift Temperature: 60C

Run Notes:



| Peak # | Start Tube | End Tube |
|---|---|---|
| 1 | A:1 | A:3 |
| 2 | A:4 | A:4 |
| 3 | A:5 | A:7 |
| 4 | A:8 | A:12 |
| 5 | A:13 | A:13 |
| 6 | A:14 | A:17 |
| 7 | A:18 | A:20 |
| 8 | A:22 | A:33 |
| 9 | A:34 | A:34 |
| --- | --- | --- |

| Duration | %B | Solvent A | Solvent B |
|---|---|---|---|
| 0.0 | 0.0 | dichloromethane | methanol |
| 2.8 | 0.0 | dichloromethane | methanol |
| 11.7 | 31.9 | dichloromethane | methanol |
| 7.5 | 50.0 | dichloromethane | methanol |
| 3.5 | 50.0 | dichloromethane | methanol |
| 1.5 | 54.8 | dichloromethane | methanol |
| 1.0 | 54.8 | dichloromethane | methanol |
| 0.8 | 55.3 | dichloromethane | methanol |
| 2.9 | 56.0 | dichloromethane | methanol |
| 1.2 | 60.1 | dichloromethane | methanol |
| --- | --- | --- | --- |

Page 1 of 1

**Figure S8**. ISCO Teledyne CombiFlash Rf+ flash chromatography purification purification of intermediate compound **4**, USC-373 after crystallization, with UV (254 nm:red trace; 280nm:purple trace) and evaporative light scattering detection (green trace). USC-373 starting eluting at 25 minutes.
